# Single Cardiac Ventricular Myosins Change Step-Size with Loading

**DOI:** 10.1101/123166

**Authors:** Yihua Wang, Chen-Ching Yuan, Katarzyna Kazmierczak, Danuta Szczesna-Cordary, Thomas P. Burghardt

**Affiliations:** Department of Biochemistry and Molecular Biology, Mayo Clinic Rochester, Rochester, MN 55905; Molecular and Cellular Pharmacology University of Miami Miller School of Medicine, Miami, FL 33136; Department of Physiology and Biomedical Engineering, Mayo Clinic Rochester, Rochester, MN 55905

## Abstract

The cardiac myosin motor powers the beating heart by catalyzed ATPase free energy conversion to contractile work. Transgenic mouse models for heart disease express mouse α-cardiac myosin heavy chain with human essential light chain (ELC) in wild type (WT), or hypertrophic cardiomyopathy linked mutant forms, A57G or E143K. Mutants modify the ELC actin binding N-terminus or C-terminus regions. Motility and single myosin mechanical characteristics show stark contrasts between the motors related to their average force, power, and displacement while all indicate the ability to down-shift ensemble step-size with increasing load. A57G and E143K consume more ATP than control WT in the presence of actin with A57G upregulating and E143K downregulating power compared with WT. Higher ATP consumption and downregulated power in E143K implies a lower unitary force. Effects on power are consistent with an A57G that impairs the ELC N-terminus actin binding and an E143K that reduces lever-arm rigidity.

## INTRODUCTION

The myosin motor protein powers muscle contraction with chemomechanical transduction of ATP free energy into the mechanical work of actin translation against resisting force. Skeletal and cardiac muscle have actomyosin interacting mostly stochastically providing power from ATP hydrolysis regulated in part with strain-dependent kinetics. Muscle myosin has a motor domain transducer containing ATP and actin binding sites, and, a mechanical coupler linking impulses from the actin bound motor to the myosin thick filament. The mechanical coupler is a lever-arm stabilized by bound light chains, essential (ELC) and regulatory (RLC), that rotates cyclically to impel bound actin. Linear actin displacement due to lever-arm rotation is the myosin unitary step-size. Myosin is a multiscaled mover producing mechanical work with single molecules translating substrate over nm distances in the cytoplasm or upscaled into highly structured ensembles with actin in the sarcomere and moving muscle tissue over distances on the order of cm.

We showed using the Qdot assay that in vitro cardiac ventricular myosin (βmys) has three distinct unitary step-sizes of ∼3, 5, and 8 nm that move actin in the absence of load with approximately 15, 50, and 35% relative step-size frequencies, respectively (1). The cardiac specific N-terminal extension of myosin ELC mediates the step-size frequency modulation (2) supporting a substantial dynamic range for velocity and force (3). Myosin RLC phosphorylation at Ser15 was shown to affect step-size frequency with unloaded phosphorylated βmys favoring 8 nm steps (4). In addition to cardiac myosin, the three step-size mechanism appears in other muscle myosins including zebrafish skeletal (5, 6).

The complexity and density of the muscle sarcomere prohibited sensing single myosin in vivo until the visible light transparent zebrafish embryo opened a window into time-resolved single myosin mechanics. The method imaged rotation or tilt of a single myosin lever-arm domain tagged with GFP in transgenic zebrafish embryo skeletal and cardiac muscle as it converts motor generated torque into the linear unitary step (5, 6). The in vivo single myosin data showed that cardiac myosin has three distinct (slightly foreshortened possibly due to strain) unitary step-sizes of ∼2, 4, and 6 nm moving actin with relative step-size frequencies that change dramatically as the muscle changes from auxotonic to near-isometric phases. These data indicate the myosin “down-shifts” average step-size by remixing the 3 divergent unitary step-sizes with changed step-frequencies. Down-shifting displacement accomplishes the myosin mechanical maneuver from a high-displacement transducer for high velocity auxotonic shortening into a low-displacement transducer maintaining tension in near-isometric contraction. The apparent correspondence between in vivo and in vitro systems implicates the ELC N-terminus in strain-dependent displacement regulation.

In the present study we introduced various constant loads on translating actin filaments and measured their effect on in vitro cardiac myosin motility and single myosin mechanical characteristics of step-size and step-frequency. We investigated the effect of load on three transgenic mouse myosins WT, A57G, and E143K. The control WT myosin has the native human cardiac ELC (MYL3) replacing mouse cardiac ELC and the mutants have the human cardiac ELC with the A57G and E143K substitutions. Pathological consequences of the hypertrophic cardiomyopathy (HCM) linked ELC mutant A57G in a transgenic mouse model indicated increases in muscle stiffness, possibly related to change in lever-arm compliance, and in myofilament calcium sensitivity leading to pathological cardiac remodeling (7, 8). In the human heart, E143K causes HCM with restrictive physiology (9). The transgenic mouse model for E143K has cardiomyopathy with enhanced passive and active tension likewise leading to pathological cardiac remodeling (10).

We find that A57G myosin mutation affects both ATPase kinetics and mechanical characteristics in contraction with higher V_max_, upregulated power, and shorter ensemble average step-size due to downregulation of the 8 nm step frequency. The latter suggests A57G impairs the actin binding function of the ELC N-terminus. E143K likewise affects ATPase kinetics and mechanical characteristics in contraction with higher V_max_ coupled to shorter unitary step-size and force that down regulates power. The N- and C-terminus localization of the A57G and E143K mutants appears to confirm the special significance of ELC N-terminus myosin generated actin movement (2) and the impact of E143K on ELC affinity for the lever-arm (11).

The WT and mutant myosins studied here, like their native in vivo counterpart in zebrafish (6), down-shift ensemble displacement under load. Step-size shifting is the adapted ensemble average step-size done by affecting step-frequency, step-size, or both, in the unitary steps. Thus the ensemble recombination of step-sizes appropriate for specific loading conditions is managed linearly by individual heads and does not necessarily require nonlinear mechanical properties acquired by the motor when integrated into the whole muscle tissue. These results confirm the longstanding observation of independent myosin impellers (12) down to the nanoscale of isolated single myosins. It also implies that the structural basis of the three step-sizes provided by the ELC N-terminus actin binding is involved in the strain-sensitivity of myosin kinetics. The latter mechanism involves strain-regulation of both the ATP dissociation of the strongly bound actomyosin complex (6) and the ADP release rate (13).

## METHODS

### Ethics

This study conforms to the Guide for the Care and Use of Laboratory Animals published by the US National Institutes of Health (NIH Publication no. 85–23, revised 2011). All protocols were approved by the Institutional Animal Care and Use Committee at the University of Miami Miller School of Medicine. The assurance number is #A-3224-01, effective November 24, 2015. Euthanasia of mice was achieved through inhalation of CO_2_ followed by cervical dislocation.

### Protein preparations

Cardiac myosin was isolated from groups of mouse hearts: WT (14 hearts), A57G (14 hearts) and E143K (6 hearts) according to Kazmierczak et al. (14). Whole hearts were isolated, the atria removed, and ventricles (left and right) were flash frozen immediately and stored at -80°C until processed. The ventricular tissue was later thawed in ice cold Guba Straub-type buffer, pH 6.5 consisting of 300 mM NaCl, 100 mM NaH_2_PO_4_, 50 mM Na_2_HPO_4_, 1mM MgCl_2_, 10 mM EDTA, 0.1% NaN_3_, 10 mM Na_4_P_2_O_7_, 1 mM DTT and protease inhibitor cocktail in a volume of 0.75 ml buffer per 0.2 g tissue. Ventricles kept on ice were first minced by hand and then homogenized for 2 minutes at 30 Hz in a Mixer-mill MM301 (Retsch). The homogenate was then incubated on ice for 40 minutes before centrifugation at 200,000 x g for 1 hour. The supernatant was then diluted 60-fold with ice cold water with 2 mM DTT and incubated on ice 30 minutes with stirring and left standing for an additional 30 minutes. The samples were centrifuged again at 8,000 x g for 10 minutes and resultant pellets were then resuspended in minimal volume of buffer containing 0.4 M KCl, 10 mM MOPS, pH 7.0, 5 mM DTT and protease inhibitor cocktail. Samples were then diluted 1:1 with glycerol, mixed gently and were shipped on ice the same day of the prep from Miami to Rochester. Proteins were stored at -20°C until used for experiments within 12 days of arrival.

### Actin-activated myosin ATPase

αmys stored in 50% glycerol was precipitated with addition of 12 volumes of ice-cold water containing 2 mM DTT, collected by centrifugation, and then resuspended in 300 mM KCl, 25 mM imidazole (pH 7.4), 4 mM MgCl_2_, 1 mM EGTA, 10 mM DTT, and 10 μg/mL leupeptin. Myosin at a final concentration of 0.6 μM was titrated with 1, 3, 5, 9, 22, 50, and 93 μM actin. The ATPase assay buffer contained 25 mM imidazole (pH 7.4), 4 mM MgCl_2_, 1 mM EGTA, 10 mM DTT, 10 μg/mL leupeptin, and a final KCl concentration of 25 mM. ATPase reaction was initiated by the addition of 3 mM ATP, and the mixture was incubated at 21 °C for 5 min. Inorganic phosphate measurements were performed using the Fiske and Subbarow method (15).

Actin-activated ATPase results were parameterized using Michaelis-Menten kinetics,

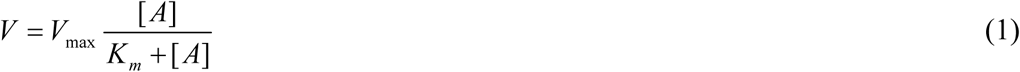

for V the ATPase rate equal to V_max_ at saturating actin concentration, actin binding constant K_m_, and actin concentration [A]. V vs [A] curves were fitted using a nonlinear algorithm to determine constants V_max_ and K_m_.

### In vitro motility and Qdot assays

*In vitro* motility and Qdot assays were performed in a flow cell using total internal reflection fluorescence (TIRF) microscopy exactly as described (2). Motility buffer included 25 mM KCl, 25 mM imidazole (pH 7.4), 4 mM MgCl_2_, 1 mM EGTA, 20 mM DTT, 10 μg/mL leupeptin, 0.7% methylcellulose, 2 mM ATP, 3 mg/mL glucose, 0.018 mg/mL catalase, and 0.1 mg/mL glucose oxidase. The flow cell was infused at the start with 0.04-0.4 μM of myosin. Actin sliding velocities for the in vitro motility assay, s_m_, and the length of actin filaments were quantitated using FIESTA software (16).

Frictional loading assays were performed as described in (17) with subtle modifications. Rhodamine labeling of actin filaments was performed with rhodamine-phalloidin and actin in a 1.2:1 molar ratio. The flow cell was infused at the start with the mixture of myosin and α-actinin in concentrations of 0.16 μM myosin and 0-6 μg/mL α-actinin (Cytoskeleton, Denver, CO).

In Qdot assays, images were acquired with an EMCCD camera (Andor, Belfast, UK) at 45 ms intervals indicated by Δt and using Andor’s SOLIS software. Each movie was recorded for 36 seconds. Intensity values were converted to photons using the conversion formula in SOLIS and the images output in TIFF format for reading into ImageJ. We tracked movement of the Qdot labeled actin at super resolution using the ImageJ plugin QuickPALM (18). All *in vitro* motility and Qdot assay experiments were conducted at room temperature (20-22 °C).

### Calibration of force in the loaded actin in vitro motility and Qdot assays

Loaded *in vitro* motility and Qdot assay data were fitted to a viscoelastic model of frictional loads as suggested by Greenberg & Moore (17). Average sliding filament velocity, s_m_, is given by,

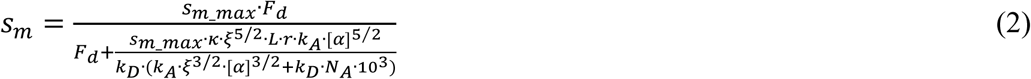

for s_m_max_ the velocity at zero load, F_d_ the ensemble myosin driving force, κ system compliance associated with α-actinin, ξ a constant defining the surface geometry of α-actinin on the flow cell surface, L actin filament length, r the reach of α-actinin to bind to actin filament, k_A_ the second-order rate constant for α-actinin attachment to the actin filament, [α] molar concentration of α-actinin, k_D_ α-actinin detachment rate, and N_A_ Avogadro’s number. s_m_max_, and F_d_ were estimated by fitting velocity data from the loaded assay using nonlinear least squares fitting in Matlab (The MathWorks, Natick, MA).

The frictional force, F_f_, exerted by α-actinin in the loaded assay is given by,

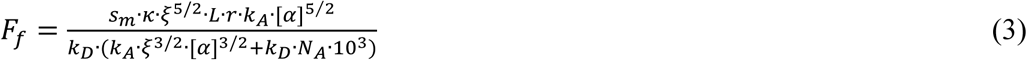

Values for parameters used in eqs. 2 & 3 are identical to those already described (17) except for fitted parameters s_m_max_, F_d_, and the length of actin filament, L ≈ 1 μm in this work. Constants in eqs. 2 & 3 (except N_A_) are estimates potentially introducing errors affecting absolute but not relative force-velocity data comparing myosins.

Frictional force-velocity curves were fit to the Hill equation (19) given by,

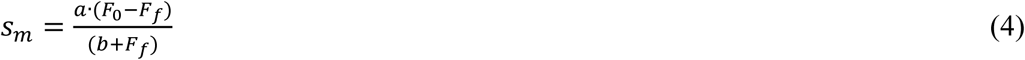

where F_0_ is the isometric force and *a* and *b* are constants. Power output, P, is given by,

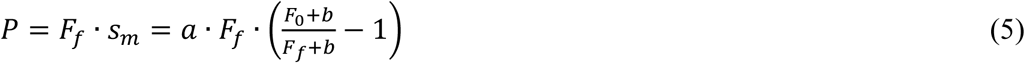

The maximal power output occurs at the point where the derivative of the power with respect to the load, F_f_, is equal to zero. The error in the power output and the load at maximal power were calculated by propagating the error from the nonlinear least squares fitting.

### Fraction of immobilized actin filaments

Average immobilized filament fraction, ω_0_, vs the frictional force is estimated from the normalized velocity,

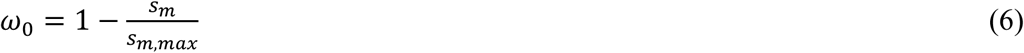

measured from loaded motility and Qdot assays.

### Qdot Assay Event-Velocity Histogram Simulation

*In vitro* motility has the myosin moving actin with a motility velocity s_m_ such that,

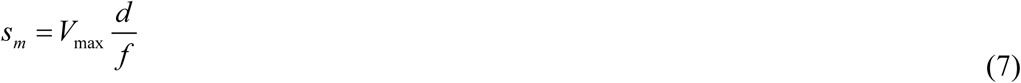

for myosin unitary step-size *d* and duty-ratio *f* (20). Duty-ratio is the time actomyosin is strongly bound during an ATPase cycle, t_on_, divided by the cycle time, 1/V_max_, hence,

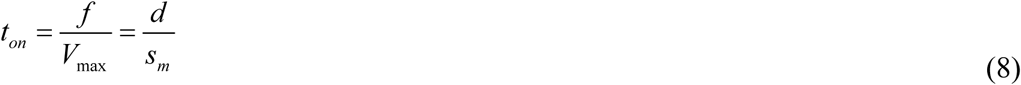

Myosin executes one of three unitary steps with step-size, d_j_, and relative step-frequency, ω_j_ for unitary step *j***=***S, I*, and *L* where S, I, and L are for the short (∼3 nm), intermediate (∼5 nm), and long (∼8 nm) nominal unitary steps of cardiac myosin (1, 4). Relative step-frequency is a characteristic proportional to the rate of cross-bridge cycling with the higher rate producing a more frequent *j*^th^ step with step-size d_j_. The dimensionless relative step-frequency is normalized with ω_s_+ω_I_+ω_L_=1. The absolute cycling rate for step *j*, V_j_, has V_j_ = V_max_ ω_j_ and with *V_max_* = ∑_*j* = *S,I,L*_.

In an ensemble of cross-bridges interacting with one actin filament, like the conditions in every muscle or motility assay, only one actin velocity is possible hence motility velocity s_m_ is the same for each unitary step-size implying each step-size has unique duty ratio and time strongly actin bound. From eq. 7, step *j* duty ratio,

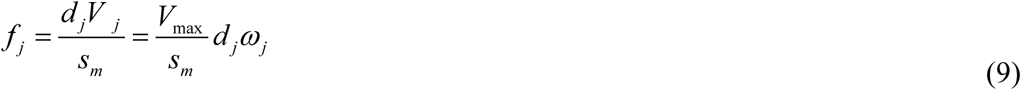

From eq. 8, the time myosin spends strongly bound to actin,

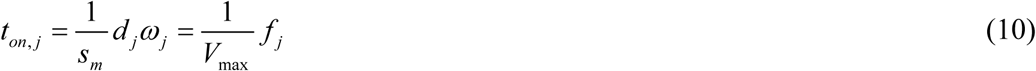

Eq. 10 affects how we simulate the event-velocity histogram since each unitary step has a unique t_on_ that varies with step-size and relative step-frequency. The t_on_ is distributed exponentially as observed using the laser trap for the in vitro actin detachment rate (21). We simulated the motility event-velocity histograms as described previously for an actin filament ∼1 μm long (2).

Ensemble average quantities used include average step-size,

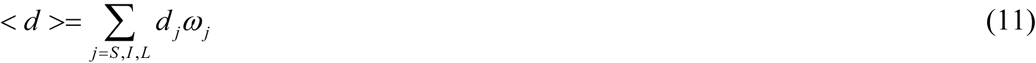

average duty-ratio <f> and time strongly actin bound <t_on_>,

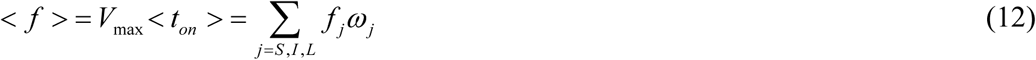

and average myosin power output from eq. 5.

V_max_ and motility velocity, s_m_, are measured under saturating actin and myosin conditions, respectively. They are constant parameter inputs to the simulation that are characteristic to each myosin tested. Simulation approximates the Qdot motility event-velocity histogram in the low velocity domain of 0∼4 natural velocity units (vu) where (d_I_/Δt) = 1 for d_I_ the intermediate step-size (near 5 nm) and where unitary events dominate. The unknown parameter set actively searched in the simulation consists of the actin binding probability for myosin (1 free parameter), step-size (3 free parameters), and relative step-frequency (3-1 = 2 free parameters due to normalization). Trial parameter values are exhaustively searched over a broad range of values set at the start of the simulation by visual inspection of the data. The simulation is programmed in Mathematica (Wolfram Research, Champaign, IL).

A Qdot assay data set consists of 8-19 acquisitions (one acquisition is one *in vitro* motility movie and corresponding event-velocity histogram) from preparations of WT, A57G, and E143K. Two or three separate protein preparations were used giving a total of 16-55 acquisitions for each protein at each actin loading. Comparison of simulated curves to data uses the χ^2^ goodness-of-fit test that is weighted by event total then summed over all the acquisitions for evaluating global goodness-of-fit.

Simulated data ensembles were created by using the 16-21 best fitting event-velocity histogram simulations generated for a Qdot assay data set. The simulations are combined linearly to approximate the measured event-velocity histogram from the pooled data (when appropriate, see below) with coefficients ≥0 while minimizing the χ^2^ goodness-of-fit test with all points equally weighted. This simulation method is identical to that described previously (2).

### Statistics

*In vitro* motility experiments using WT, A57G, and E143K preparations corresponded to 16-55 independent event-velocity histograms for each protein. We simulated data from each protein preparation independently to estimate single myosin mechanical characteristics consisting of 3 step-sizes and 3 step-frequencies. We compared 3 step-sizes or 3 step-frequencies as categorical variables in factor 1 and the 2-3 independent data sets acquired for the separate protein preparations (with 8-19 independent event-velocity histograms for each species) in factor 2 using two-way ANOVA with Bonferroni or Tukey-Kramer post-tests for significance at the 0.05 level. This test indicated no significant difference among the independent data sets for each species hence data sets were pooled for a given protein.

## RESULTS

### Expression levels of human ELC

**Figure 1** shows the SYPRO Ruby stained SDS-PAGE gels for mouse αmys species. NTg is the nontransgenic mouse myosin. Staining intensity for native RLC and ELC compared to the human ELC species WT, A57G, and E143K indicate 79% human native ELC, 75% human A57G, and 58% human E143K with the remainder the native mouse isoform.

**Figure 1.**
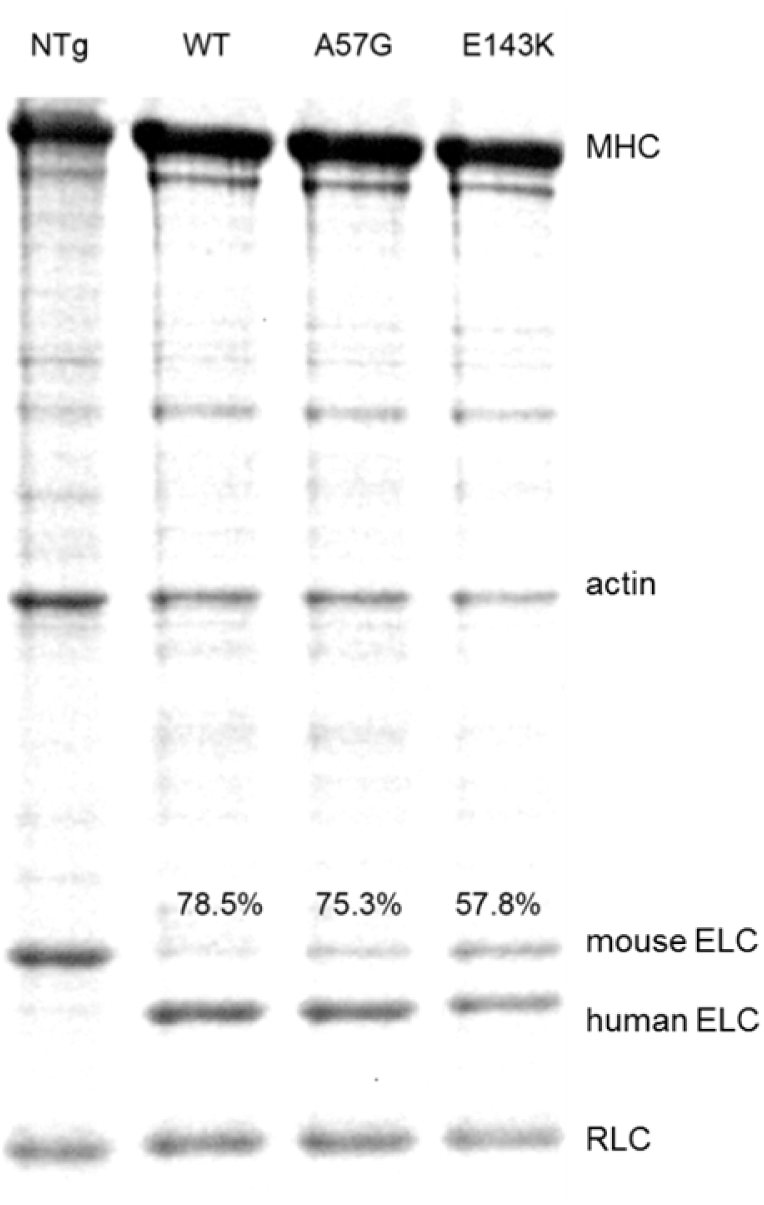
SDS-PAGE of nontransgenic (NTg) and transgenic WT, A57G, and E143K myosin preparations. MHC is the mouse cardiac α-myosin heavy chain. Human vs total ELC is indicated with %.

### Actin-activated ATPase of cardiac myosin

**Figure 2** shows the actin-activated ATPase vs actin concentration, [A], for WT (red circles), A57G (blue triangles), and E143K (black squares) with error bars indicating standard deviation. Actin-activated myosin ATPase on WT, A57G, and E143K distinguished each protein species mainly by their V_max_ with A57G and E143K nearly 2× higher than WT. **Table 1** summarizes the Michaelis-Menten parameters and the sampling statistics for the data in **Figure 2**.

**Figure 2.**
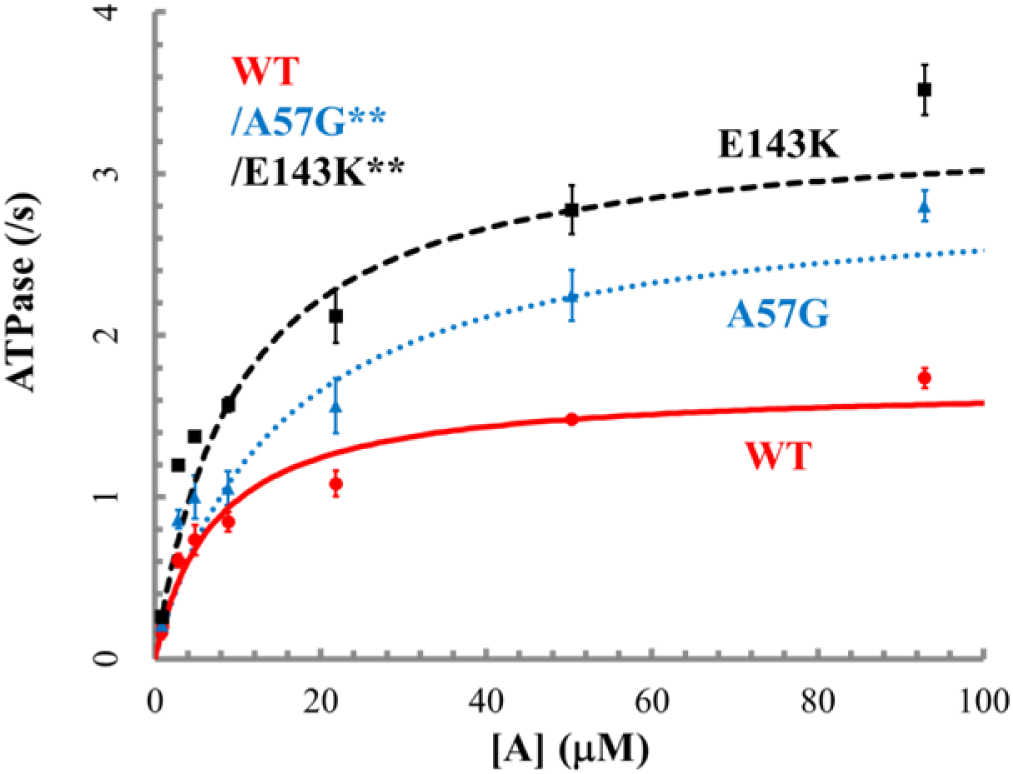
Actin-activated myosin ATPase for wild type (WT) and mutants E143K and A57G expressed in transgenic mouse ventriculum. [A] is actin concentration. Error bars show standard deviation for sampling statistics given in Table 1. Conditions are given in Methods. Significance of ATPase rate vs [A] data in pairwise comparison for the WT and mutant species is indicated. They differ significantly with confidence level p < 0.01 as indicated by **\*\***

**Table 1:** Average mechanical characteristics of transgenic myosins^1^.

| **Table 1: Average mechanical characteristics of transgenic myosins<sup>1</sup>** |  |  |  |  |
| --- | --- | --- | --- | --- |
|  | WT | A57G | E143K | p< |
| Actin-activated myosin ATPase |  |  |  |  |
| $V_{\max}$ (1/s) | $1.69 \pm 0.06$ (n=6) | $2.90 \pm 0.16$ (4) | $3.32 \pm 0.01$ (2) | 0.01 |
| $K_m$ (μM) | $7.22 \pm 0.54$ | $15.0 \pm 3.21$ | $9.81 \pm 0.49$ | - |
| Peak in vitro motility velocity for unloaded actin |  |  |  |  |
| $s_m$ (μm/s) | $0.81 \pm 0.05$ (n=32) | $0.91 \pm 0.05$ (38) | $0.74 \pm 0.05$ (15) | 0.01 |
| Isometric force in pN |  |  |  |  |
| $F_{\text{iso}}$ (eq. 2) | $514.7 \pm 34.7$ (n=11) | $699.0 \pm 52.4$ (7) | $373.0 \pm 37.4$ (7) | 0.01 |
| $F_{\text{iso}}$ (eq. 4) | $308.8 \pm 16.4$ | $344.5 \pm 20.6$ | $245.5 \pm 23.3$ | 0.01 |
| Peak power @ load |  |  |  |  |
| $P_{\max}$ (fW) | $0.049 \pm 0.005$ | $0.060 \pm 0.007$ | $0.032 \pm 0.006$ | NA |
| @ load (pN) | $141.0 \pm 6.9$ | $155.8 \pm 8.5$ | $114.0 \pm 10.1$ | NA |

<sup>1</sup>Average values and standard deviation indicated for (n) replicates. ANOVA significance p< indicates confidence level for distinguishing WT/A57G and WT/E143K pairs. NA indicates not applicable for error derived from fitting eq. 4 and using eq. 5.

We tested significance of data in **Figure 2** using 2-way ANOVA with factor 1 proteins WT, A57G, and E143K, and factor 2 the actin concentration [A]. We find the data sets differ significantly with confidence levels p < 0.01 (**) in pairwise comparisons of WT with mutants. The HCM implicated ELC mutants substantially impact myosin kinetics by speeding up actin-activated hydrolysis without affecting weak actin binding constant K_m_.

### In vitro motility

**Figure 3** shows motility velocity, *s_m_*, vs myosin bulk concentration, [M], for WT (red circles), A57G (blue triangles), and E143K (black squares) with error bars indicating standard deviation. Smooth lines are the sum of two fitted exponential curves to indicate trends. Motility velocity increases with increasing αmys bulk concentration until reaching maximum at 0.14-0.20 μM and then slightly decreasing. Peak motility velocities and sampling statistics are given in **Table 1**. The WT motility velocity vs [M] compares favorably with our earlier measurement of this relationship (2).

**Figure 3.**
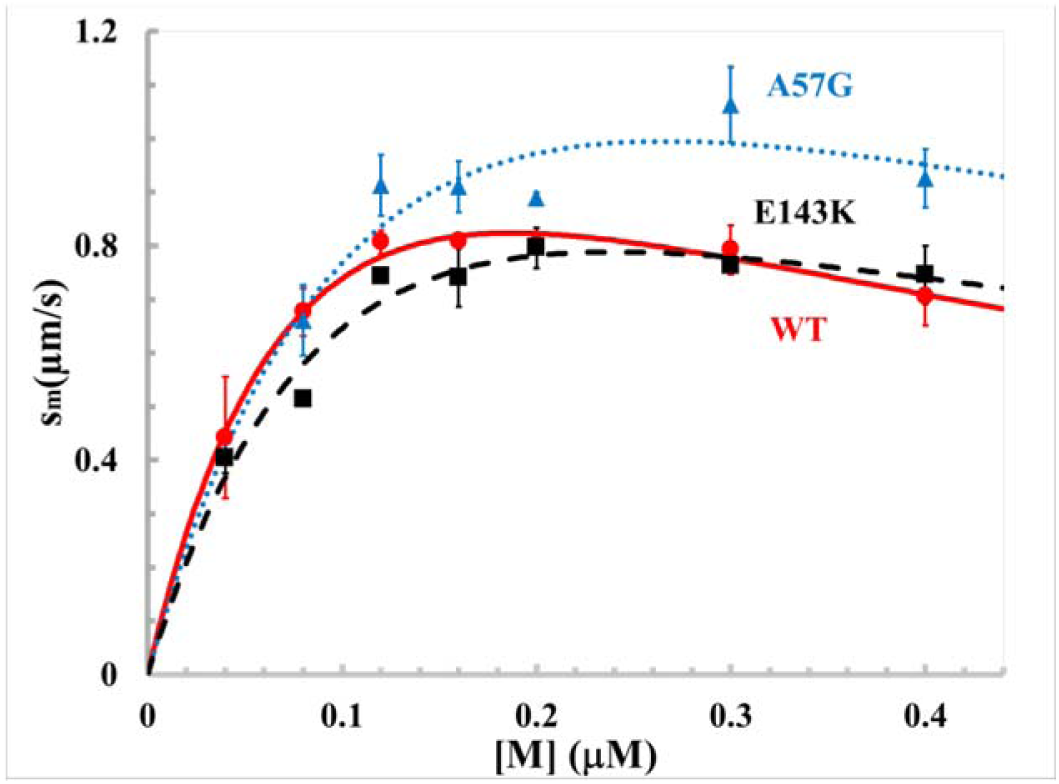
Motility velocity, s_m_, for the myosin isoforms moving rhodamine labeled actin. Error bars show standard deviation for sampling statistics given in Table 1.

### In vitro force measurements show impact of mutation

**Figure 4** shows motility velocity, *s_m_*, vs α-actinin concentration (**panel a**) or loading frictional force, F_f_ (**panel b**), exerted by α-actinin in piconewtons (pN) for WT (red circles), A57G (blue triangles), and E143K (black squares). The α-actinin concentration was converted to a calibrated frictional force using eq. 3. Fitted curves are based on eqs. 2 and 4 (Hill equation) for **panels a & b**. Error bars in **Figure 4** indicate standard deviation for sampling statistics given in the caption. The isometric force obtained from fitting the data in **Figure 4** is summarized in **Table 1**.

**Figure 4.**
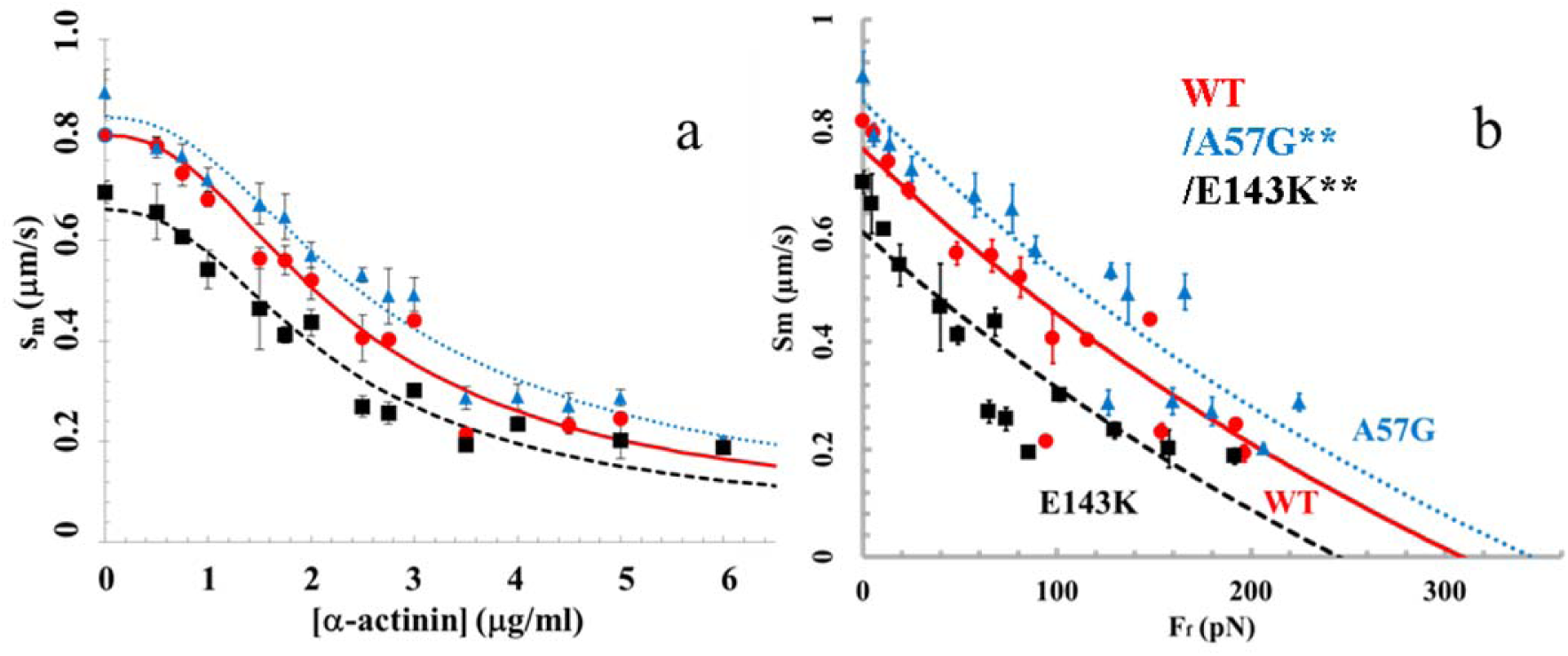
Motility velocity, s_m_, for myosin isoforms WT (red circles), A57G (blue triangles), and E143K (black squares) vs α-actinin concentration (**panel a**) or calibrated frictional force, F_f_, exerted by the α-actinin (**panel b**). Sample sizes for the standard deviations indicated by error bars are WT (n=11), A57G (n=7), and E143K (n=7) for n the number of acquisitions corresponding to 1 in vitro motility movie per acquisition. s_m_ vs F_f_ data in pairwise comparison for the WT and mutant species is indicated for each pair in **panel b**. They differ significantly with confidence level p < 0.01 as indicated by **.

We tested significance of data in **Figure 4 panel b** using 2-way ANOVA with factor 1 proteins WT, A57G, and E143K, and factor 2 the frictional force F_f_. We find the curves differ significantly for each pairwise comparison of WT with mutants and with confidence levels p < 0.01 (**).

**Figure 5** shows power (s_m_ × F_f_ from **Figure 4**) vs F_f_ for WT (red circles), A57G (blue triangles), and E143K (black squares) with error bars indicating standard deviation for sampling statistics given in the caption. Power is in units of femtowatts (fW). Fitted curves are based on eq. 5. Peak power and the load at peak power are summarized in **Table 1**.

**Figure 5.**
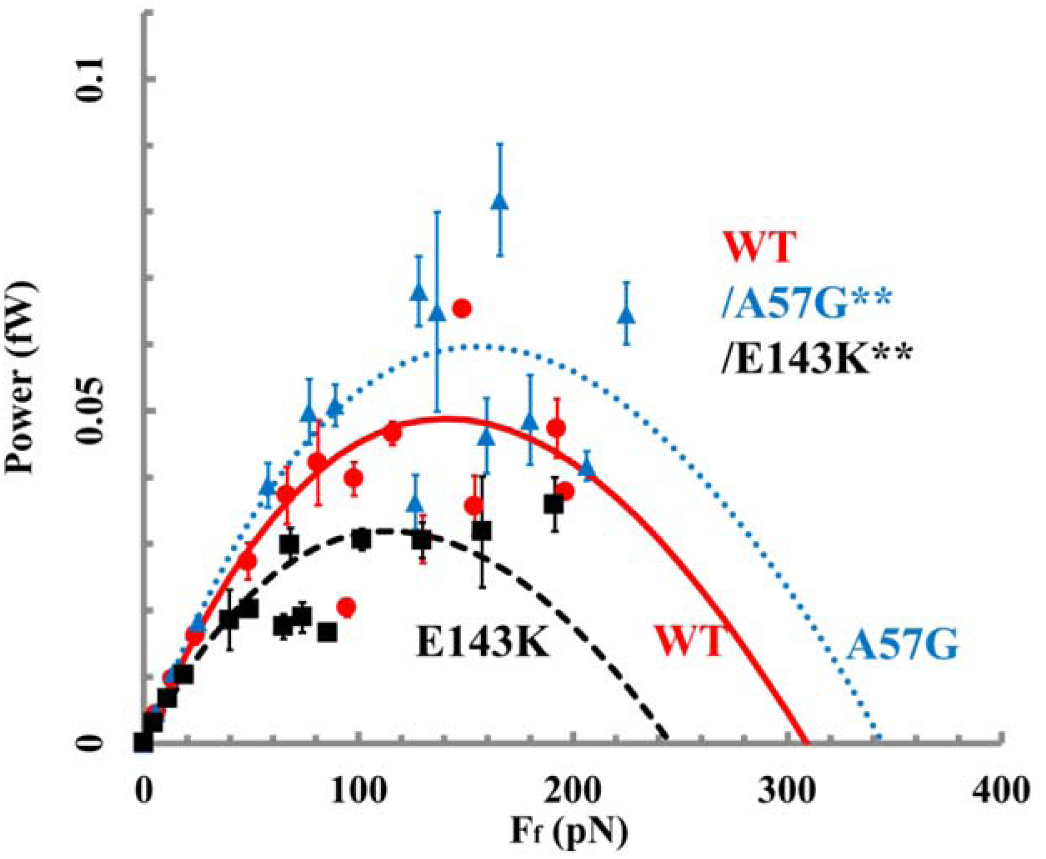
Power produced by WT (red circles), A57G (blue triangles), and E143K (black squares) myosin isoforms in the motility assay vs F_f_, the frictional force exerted by α-actinin. Sample sizes for the standard deviations indicated by error bars are WT (n=11), A57G (n=7), and E143K (n=7). Power vs F_f_ data in pairwise comparison for the WT and mutant species is indicated for each pair. They differ significantly with confidence level p < 0.01 as indicated by **.

We tested significance of data in **Figure 5** using 2-way ANOVA with factor 1 proteins WT, A57G, and E143K, and factor 2 the frictional force F_f_. We find the curves differ significantly for each pairwise comparison of WT with mutants and with confidence levels p < 0.01 (**).

The HCM implicated ELC mutants substantially and differently change the *in vitro* force-velocity characteristics of the cardiac myosin with A57G enhancing, and E143K diminishing, power generated compared with WT.

### Loaded Qdot assay models the working muscle

The Qdot assay has labeled actin filaments ∼1 μm long translating over surface bound mouse αmys. **Figure 6** shows the Qdot assay event-velocity histogram pooled data from 16-55 acquisitions for each species (WT, A57G, and E143K) and for increasing load as indicated in pN. Columns are nearly equal frictional loads for the rows with identical transgenic myosins. The event-velocity histograms cover the low velocity domain of 0∼4 natural velocity units (vu) where (d_I_/Δt) = 1 for d_I_ the intermediate step-size of ∼ 5-6 nm and frame capture interval Δt = 45 ms. Summary data (solid squares) has the baseline contribution from thermal/mechanical fluctuations (red solid line) already subtracted to give motility due to myosin activity. Peaks or inflection points appearing below 2 vu are short (short red up arrow or S), intermediate (d_I_ and longer green down arrow or I), and long (longest blue up arrow or L) step-sizes in nm. Green and blue arrows also indicate a few of the unitary step combinations. Step-sizes are indicated by the color-coded numbers nearest the arrows. Step-size significance was tested using 1-way ANOVA comparing WT, A57G, and E143K species. The intermediate step-size for the E143K species under 0 load differs significantly from the equivalent WT species with confidence level p < 0.01 as indicated by ** in **Figure 6**. The significance suggests a shorter intermediate and average step-size for E143K compared to WT.

**Figure 6.**
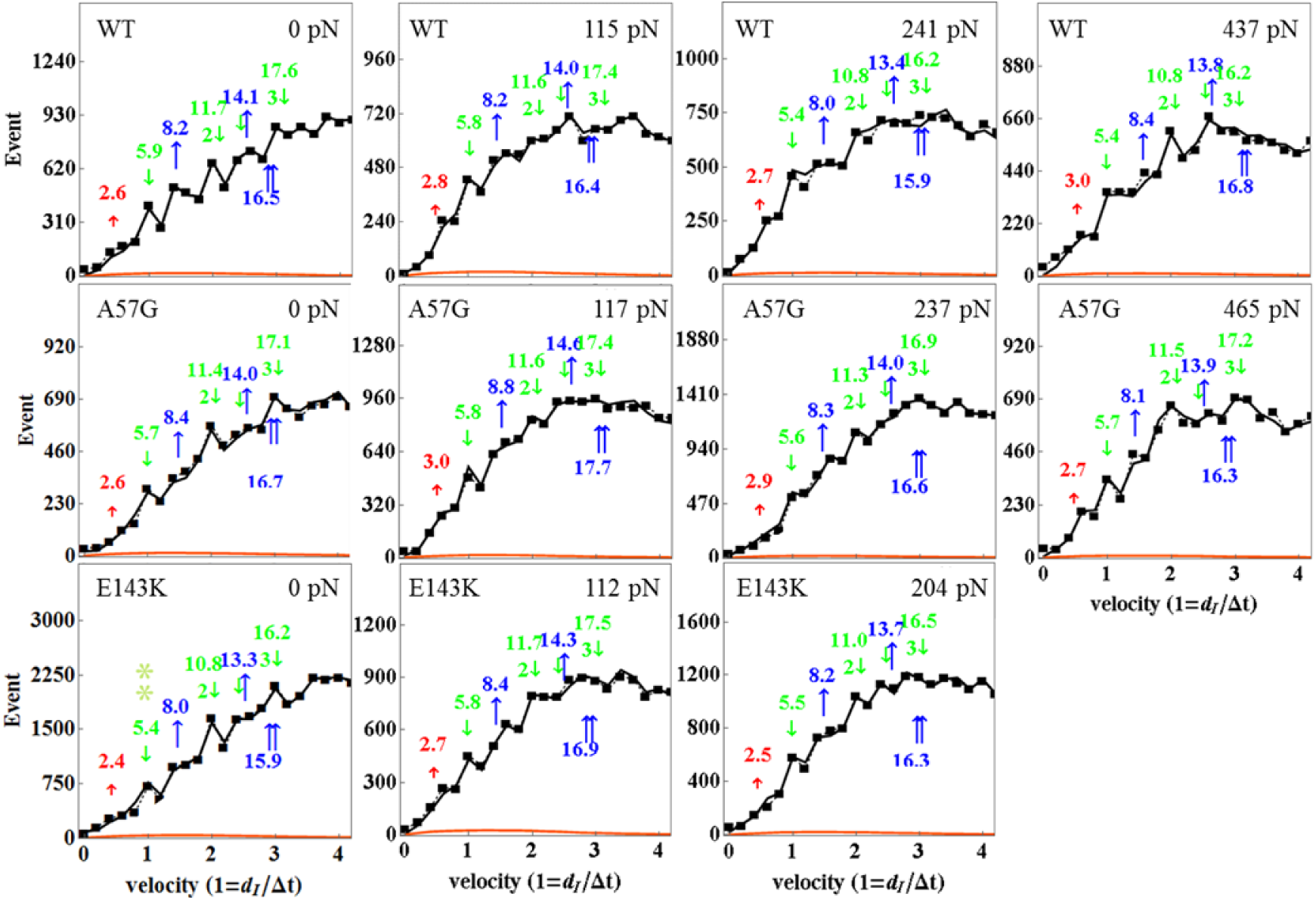
Event vs velocity histograms (solid squares) for WT, A57G, and E143K myosins and for frictional actin loading, F_f_, from 0 to 465 pN. The solid red line is the baseline due to thermal/mechanical fluctuations that was subtracted from the raw data to give the data points at solid squares. Black lines are simulations conducted as described in Methods and used to estimate step-size (at arrows) and step-frequency (see **Figure 7**). Natural velocity units (vu) have (d_I_/Δt) = 1 for d_I_ the intermediate step-size (green down arrow at ∼ 5-6 nm). Step-sizes have standard deviation of ∼0.5 nm for replicates described in Methods under *Statistics*. The intermediate step-size for the E143K species under 0 load differs significantly from equivalent WT species with confidence level p < 0.01 as indicated by **. Other step-sizes are not significantly different for confidence level p < 0.05.

**Figure 7** shows the step-frequency (ω) expectations, expectation values, and standard deviations for the S, I, and L unitary steps derived from each event-velocity histogram in a one-to-one relationship with the panels in **Figure 6**. The expectation curves indicate the relative probability for step-frequency along the abscissa. The area under the color coded curves for the S, I, and L steps equals expectation values ω_S,_ ω_I_ and ω_L_ respectively. The sum ω_S_ + ω_I_ + ω_L_ = 1 in each panel.

**Figure 7.**
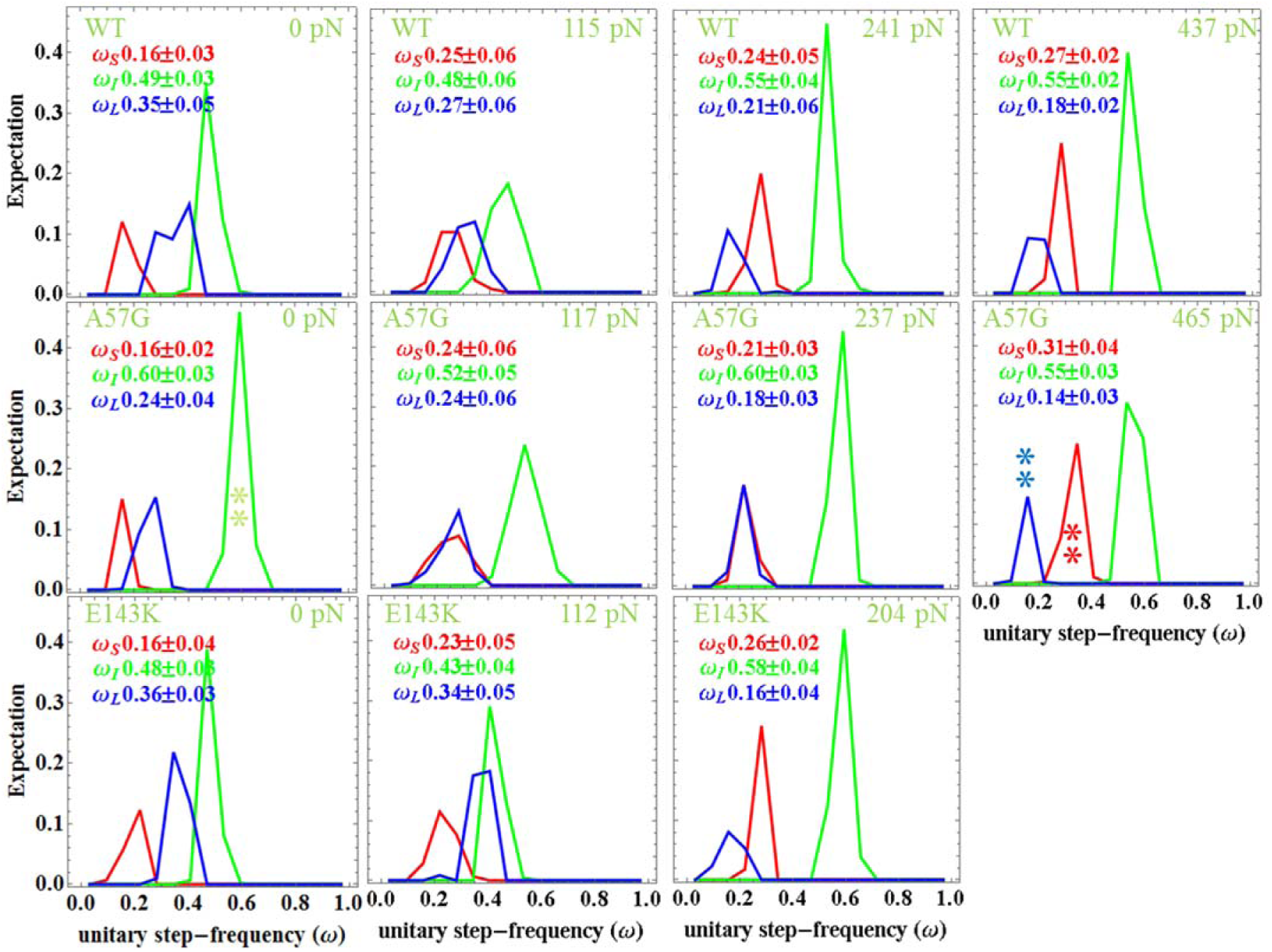
The step-frequencies for the WT (top row), A57G (middle row), and E143K (bottom row) under frictional loads indicated in each panel in pN units. Columns are equivalently loaded actin for each transgenic protein. Errors are standard deviations for replicates described in Methods under *Statistics* Intermediate step-frequency for the A57G species differs significantly from equivalent WT species with confidence level p < 0.01 for no load and as indicated by **. Other step-frequencies for the A57G and WT species also differ significantly with confidence level p < 0.01 under maximum load and as indicated by **.

Step-frequencies in **Figure 7** for F_f_ = 0 indicate WT and E143K are statistically identical while A57G favors the 5 nm step with a step-frequency of ∼60% compared to ∼50% in WT and E143K. This difference is significant at confidence level p < 0.01 (indicated by **). The canonical step-frequencies of 13, 50, and 37% observed in WT and E143K (and βmys) shifts in favor of the 5 nm step in A57G. We attribute this to the perturbation of the actin binding ELC N-terminus by the mutation at A57. Earlier work demonstrated that the step-frequencies for the 5 and 8 nm steps were similarly modulated by truncation of the ELC N-terminus (2). Direction and amplitude of the step-frequency adjustment for the truncated ELC is identical to within error to that shown in **Figure 7** for A57G.

Step-frequencies in **Figure 7** indicate the 5 nm step-size is always predominant, however, at the highest loads the second most frequent step-size reverses from 8 to 3 nm repeating a down-shifting maneuver first observed in vivo from cardiac myosin in zebrafish embryo hearts (6). Step-frequencies for the 3 and 8 nm step-sizes are significantly different from each other in WT, A57G, and E143K at a confidence level of p < 0.01. Cardiac myosin replaces 5 with the 3 nm step as the predominant step at isometric contraction in the zebrafish heart. This does not happen in vitro possibly because loading is less when α-actinin imposes the drag on actin movement (17).

The WT and mutant myosins differ on how they accomplish the 8 to 3 nm step-size probability reversal with A57G requiring a higher loading force than WT or E143K, however, all species down-shift over the loading conditions tested. The overall effect of the step-size shifting for each protein is to shorten the ensemble average step-size, <d>, as load increases and as indicated in **Figure 8 panel a**.

**Figure 8.**
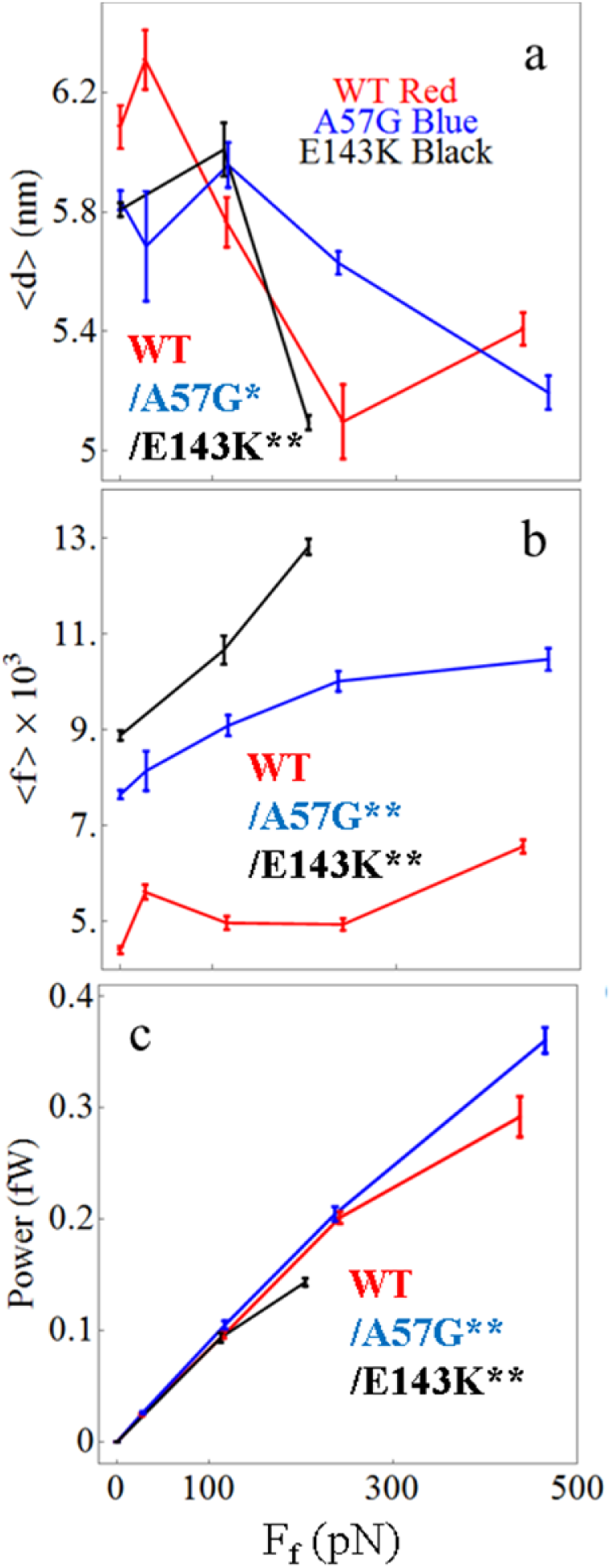
In vitro single myosin average step-size <d> (**panel a**), average duty ratio (**panel b**), and power (**panel c**) for WT (red), A57G (blue), and E143K (black) αcardiac myosin measured using the Qdot assay with retarding force. Errors are standard error of the mean for the 16-55 replicates described in Methods under *Statistics*. Data in pairwise comparison for the WT and mutant species is indicated for each panel on the right. They differ significantly with confidence level p < 0.05 or 0.01 indicated by * or **.

**Figure 8** indicates the ensemble averaged quantities for the WT, A57G, and E143K species computed using eqs. 10 & 11 from the single myosin characteristics summarized in **Figures 6 & 7. Figure 8 panel b** shows duty-ratio vs load. Control WT has a lower and flat duty ratio with increasing load implying that the fraction of force producing myosins during contraction at these loads is low and static. Compared to WT, A57G and even more so E143K, have high and rising duty ratio with increasing load implying that the fraction of force producing myosins is higher and growing during the contraction phase. **Figure 8 panel c** shows larger power production for A57G indicating hypercontractility. Myosin hypercontractility associates with HCM disease (22) but for reasons that probably also include secondary effects induced by heart tissue remodeling (8). For the same reasoning, hypocontractility in E143K (again **panel c**) is unexpected and suggesting E143K unitary force is compromised given that it has higher duty ratio but produces less work. This is possible if E143K reduces lever-arm rigidity. Overall ELC positioning of A57G and E143K seems to fit this scenario with A57G directly impacting the ELC N-terminus actin binding during the active cycle and E143K passively reducing lever-arm rigidity.

We tested significance of data in **Figure 8** using 2-way ANOVA with factor 1 proteins WT, A57G, and E143K, and factor 2 the frictional force F_f_. We find that these data differ significantly for each pairwise comparison of WT with mutants and with confidence levels p < 0.01 (**) or p < 0.05 (*).

Step-size modulation for the E143K mutant (**Figure 6**) and step-frequency modulation for the A57G mutant (**Figure 7**) are mechanical manifestations of the mutations detected in the unloaded conditions. They cause <d> to shorten either directly by lowering the intermediate step-size in E143K or by the failure of the ELC N-terminus to bind actin in a timely manner and so frequently failing to complete the 8 nm step-size in A57G (see zero load in **Figure 8 panel a**).

Step-frequency modulation is the signature characteristic of the ELC N-terminus actin binding interaction (2). Recent work on the in vivo zebrafish heart suggests the ELC N-terminus possesses strain-sensing characteristics when actin bound that has a ratchet-like selective resistance to movement in the direction of the loading force while permissive to movement in the direction of the contractile force. Consequently, the step-frequency modulation caused by the loading of the actin filament in the myosins tested here (**Figure 7**) implies a role for the ELC N-terminus actin binding in the response of single myosins to load. Here, it causes <d> to shorten as load increases by up modulating shorter 3 & 5 nm step-sizes at the expense of the longer 8 nm step-size. For the in vivo zebrafish heart application we introduced a 4-pathway model for the myosin contraction cycle that rationalizes the effect of load on <d> (6). In the next section we use Qdot assay data in the 4-pathway model illustrating the close correlation between in vitro and in vivo contexts of myosin based contraction.

### Contraction cycle 4-pathway model

Work using the unloaded in vitro Qdot assay on βmys (1) combined with the in vivo single cardiac myosin imaging of zebrafish embryo hearts (6) suggested a 4 pathway network for generating 8, 5, 5+3, and 3 nm myosin step-sizes (**Figure 9**). Earliest work using the unloaded Qdot assay alone identified the 8 (blue path), 5 (green path lower branch), and 5+3 (green path upper branch) nm step-size pathways (1). The vivo system added the context of the intact auxotonic and near-isometric muscle that identified the solo 3 nm step-size (red) pathway (6).

**Figure 9.**
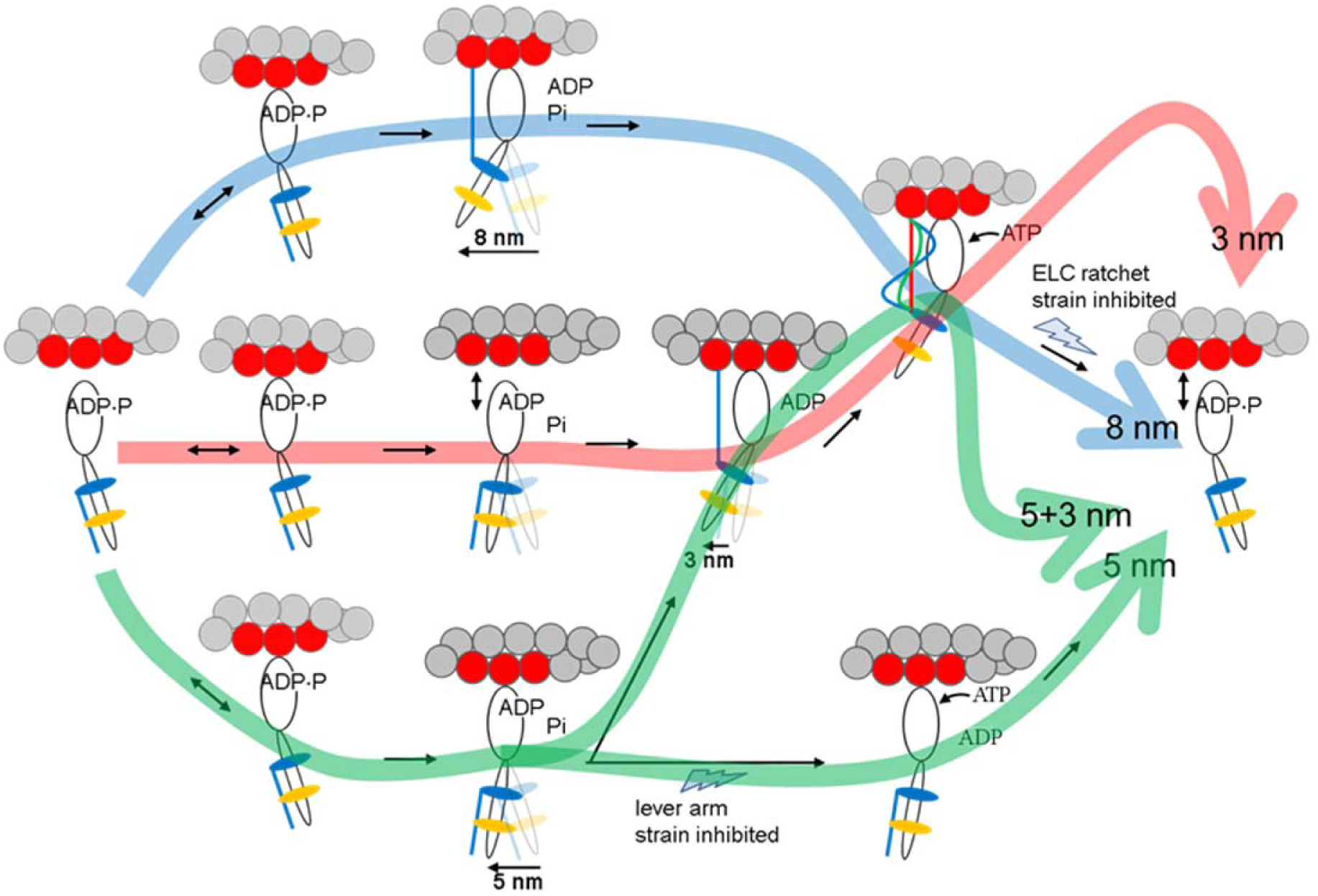
The 4 pathway network for in vivo and in vitro cardiac myosin unitary steps.

In **Figure 9**, weak actin binding myosin reversibility (indicated with ↔) preemptively avoids pathways that do not complete a cycle under increasing load. Under loaded conditions, flux through the cycle is checked at two strain inhibited points. The traditional lever arm strain inhibited checkpoint when under strain inhibits ATP dissociation of myosin following the 5 nm step-size (green path lower branch) instead sending flux towards the 3 nm step-size (green path upper branch). The new ELC-ratchet checkpoint inhibits the 3, 8, and 5+3 nm pathways (red, blue, and upper branch green). The latter senses tension in the actin bound ELC N-terminus such that the slack ELC extension (blue, when muscle is unloaded or rapidly shortening under low load) detaches quickly to allow completion of the cycle, the moderately tense ELC extension (green, when muscle is under moderate loads approaching isometric) detaches slower to exert static myosin tension contributing to force on slowly translating muscle filaments, and the maximally tense ELC extension (red, when muscle is near isometric) detaches slowest to exert static myosin tension contributing more of the force on the static muscle filaments. Strongly bound myosins contributing static tension are the force bearing 0 length step-size myosins not explicitly depicted in **Figure 9** (rather see **Figure 8** in (6)). Modulating flux through 2 strain-dependent steps with different inhibit efficiencies adjusts average step-size. The favored pathway at high tension involves the 3 nm step implying the ELC ratchet strain based checkpoint is less inhibiting than the lever arm strain checkpoint.

We described how to compute the myosin flux through the 4 pathway network in **Figure 9**. In addition to the step-frequencies provided in **Figure 7**, we must also estimate the force bearing 0 length step-size step-frequency, ω_0_, quantitating myosins contributing static tension. The ω_0_ step-frequency rises with load and estimates the immobilized fraction of Qdot labeled actin filaments using eq. 6. **Figure 10** shows the immobilized fraction data for the WT, A57G, and E143K species plotted over the frictional drag forces imposed by α-actinin. These data are also indicated in **Table 2**. Step-sizes depicted graphically in **Figure 9** have their step-frequencies indicated in **Table 2**. All myosin species down-shift ensemble displacement with increasing resistive load by remixing its 3 divergent unitary step-sizes with changed step-frequencies in a mechanism paralleling that observed in vivo.

**Figure 10.**
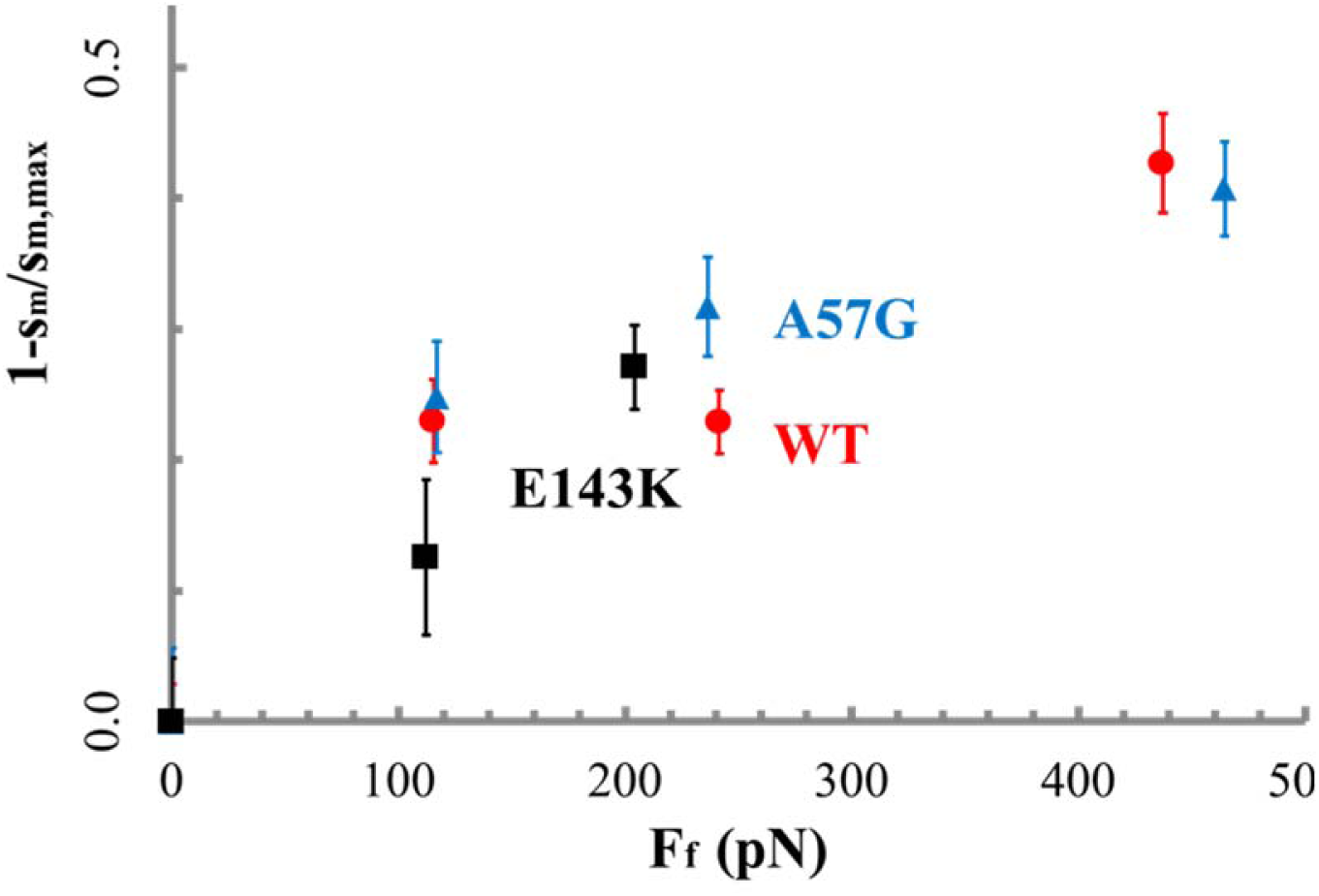
The immobilized actin filament fraction vs loading force estimated from the motility of Qdot labeled actin and eq. 6. Errors are standard deviation for the 16-55 replicates described in Methods under *Statistics*.

**Table 2:** Myosin flux through the 4-pathway network.^1^.

| **Table 2:** Myosin flux through the 4-pathway network.<sup>1</sup> |  |  |  |  |  |
| --- | --- | --- | --- | --- | --- |
|  | WT |  |  |  |  |
| $F_r$ (pN) | $\omega_0$ | 8 nm | 5 nm | solo 3 nm | 5 + 3 nm |
| 0 | 0 | 39±4 | 43±4 | 6±3 | 12±3 |
| 115 | 0.23±0.06 | 29±8 | 43±10 | 16±10 | 12±5 |
| 241 | 0.23±0.05 | 24±7 | 49±10 | 14±9 | 13±5 |
| 437 | 0.43±0.08 | 19±5 | 48±12 | 24±11 | 9±5 |
| p< |  | 0.01 | - | 0.01 | - |
|  | A57G |  |  |  |  |
| 0 | 0 | 27±3 | 54±4 | 7±2 | 12±3 |
| 117 | 0.25±0.09 | 26±9 | 45±11 | 17±11 | 12±5 |
| 237 | 0.32±0.08 | 21±5 | 52±13 | 15±11 | 12±5 |
| 465 | 0.41±0.07 | 14±5 | 50±13 | 27±13 | 9±5 |
| p< |  | 0.01 | - | 0.01 | - |
|  | E143K |  |  |  |  |
| 0 | 0 | 40±2 | 42±4 | 7±2 | 11±3 |
| 112 | 0.13±0.11 | 37±8 | 37±11 | 17±12 | 9±4 |
| 204 | 0.27±0.06 | 17±6 | 52±13 | 18±12 | 13±5 |
| p< |  | 0.01 | - | 0.01 | - |

<sup>1</sup>Average values indicated with standard deviation for the 16-55 replicates described in Methods under *Statistics*. Normalization has the sum of step-frequencies for the four step-size categories equal to 100%. Significance values in the p< rows are for the WT, A57G and E143K species and indicating the 1-way ANOVA comparison of step-frequency for 0 and maximum load cases. p<0.01 implies the step-size column has step-frequencies that differ due to load while – indicates no significant difference. Pathways for the 8, 5, solo 3, and 5 + 3 nm step-sizes are depicted graphically in **Figure 9**.

## DISCUSSION

The high-throughput Qdot assay has myosin immobilized on a planar glass substrate and measures motor step-size in the context of an ensemble of actomyosin interactions. The native myosin impels Qdot labeled actin with nanometer scale actin displacement measured using super-resolution microscopy (23) under prismless total internal reflection illumination (24). The assay has negligible compliance to faithfully mechanically characterize the isolated myosin motor (2). Prior work with unloaded actin in the Qdot assay indicated cardiac myosin moves actin with three distinct unitary step-sizes and with characteristic step-frequencies (1). Here we introduce various constant loads on the translating actin filaments and measure their effect on motility, myosin step-size, and myosin step-frequency. In vitro loading emulates the cardiac ventriculum where a heartbeat circumscribes changing modes from relaxation, to auxotonic shortening, to isometric contraction.

Zebrafish embryo heart in vivo single myosin mechanical data indicates cardiac myosin down-shifts average displacement by remixing the 3 unitary step-sizes with changed step-frequencies (6). Down-shifting occurs on the fly from a high-displacement/low-force transducer for high velocity auxotonic shortening into a low-displacement/high-force transducer maintaining tension in near-isometric contraction. We report here qualitatively equivalent myosin down-shifting in vitro using the loaded Qdot assay. Data in **Figure 7** shows the step-frequencies for transgenic mouse αmys with human ELC (WT) in the top row with loading force from α-actinin increasing from left to right. Increasing load remixes contributions from the longest and shortest step-sizes to reduce the ensemble average step-size, <d>, shown in **Figure 8**. Ensemble averaged duty-ratio and power also in **Figure 8** indicate other critical systemic adaptations to the loaded system by ensemble myosin resulting from the single myosin step-frequency adaptation indicated in **Figure 7**. Overall, our data demonstrates that the recombination of step-size frequencies for specific loaded conditions adapts the myosin motor to a changing load demand in an evident correspondence with the in vivo system. It implies that the varied combination of step-sizes for specific loaded conditions, first observed in vivo, does not require hierarchical mechanical properties acquired by motor integration into whole muscle. It also implies, like the in vivo results (6), that the structural basis of the three step-sizes provided by the ELC ratchet is also involved in the myosin strain-sensitivity.

We address the basis of familial heart disease and significance of the ELC ratchet with the two ELC mutants, A57G and E143K. They cause distinct disease phenotypes, occupy opposite ends of the myosin light chain sequence, and differently affect single αmys mechanics. The A57G and E143K mutants cause hypertrophy in human hearts while E143K is further differentiated as causing restrictive physiology. In vitro, myosins containing the A57G or E143K mutations have ∼2× larger V_max_ than control WT but unaffected K_m_ in actin-activated myosin ATPase. Motility velocity is little affected (**Table 1**). Unique differences between A57G and E143K emerge in motility data without and with the imposition of a load on the actin filaments. Calibrated force/velocity curves in **Figure 4** imply power up regulation in A57G and down regulation in E143K. This effect is confirmed by the power vs force curves in **Figure 5** for WT, A57G, and E143K.

In the unloaded Qdot assay the A57G and E143K mutants stand out due to their altered step-frequencies and step-size compared to control WT, respectively. Step-frequencies for A57G resemble those observed from N-terminal truncated ELC myosins in porcine or mouse cardiac myosin (2) and imply A57G disables the ELC ratchet. The intermediate step size of ∼5 nm for E143K is significantly shortened compared to control. We also noted that <d> for the unloaded mutants are both less that for WT (**Figure 8 panel a**). For A57G it is due to step-frequency modulation by the mutation (**Figure 7** top row). For E143K it is due to the significant step-size reduction (**Figure 6** top row). We suggested E143K myosin produces a lower unitary force following analysis of the full (loaded and unloaded) Qdot assay data set. Consistent with all available data is the implication that E143K passively reduces lever-arm rigidity. The unloaded Qdot assay data, indicating the shorter intermediate step-size for E143K, evidentially also implies this possibility.

The imposition of load on the actin filament is reflected in the step-frequencies for each protein species in **Figure 7**. WT, A57G, and E143K all respond distinctively to the increasing load however at the highest loading force they similarly favor the 3 nm over the 8 nm step. The myosin flux through the 4 pathway network in **Figure 9** gives a more detailed accounting for the step-size adaptation to load that tallies an expanded set of step-size frequencies for each myosin species and loading condition. The tally uses the step-frequency information in **Figure 7** and an estimate for the immobilized fraction of actin filaments that rises with load according to the data in **Figure 10. Table 2** summarizes flux through the system in 4 step-size categories that separate probability for the unique solo 3 nm step-size from the 3 nm step-size coupled to its antecedent 5 nm step. The tallies show significant incremental decreases in the 8 and increases in the solitary 3 nm step-frequencies while 5 nm and 5+3 nm step-frequencies are about the same.

Performance contrast among the three myosin species is evident in the ensemble average data (**Figure 8**). Duty ratio vs load for the A57G is displaced upward compared to WT by the constant and larger V_max_ while curve shapes differ from their contrasting dependence on step-frequency vs load (see eqs. 9 & 12). The effect of mutation on duty ratio vs load is similar but amplified in E143K due to the larger change in V_max_. A57G and E143K have high and rising duty ratio with increasing load compared to WT implying their ensemble will have more force producing myosins. If actively cycling vs super-relaxed (25) myosins re-equilibrate in the sarcomere during the phases of a heartbeat, the shift to strongly actin bound myosins implied by the duty ratio has the ELC mutants recruiting more super-relaxed myosins to the active state during the contraction phase than the native myosins. Either way, more ATP consumption will impact health in the long term.

Average power (**Figures 5 & 8 panel c**) up regulation in A57G is expected with the higher ATP consumption and its association with HCM (22). E143 is also associated with HCM but average power down regulation is observed. This is expected if E143K reduces lever-arm rigidity. Overall positioning of substitutions A57G and E143K in ELC seems to fit this scenario with A57G in the N-terminus directly impacting actin binding during the active cycle and E143K passively reducing lever-arm rigidity probably because of weaker E143K binding to myosin (11). A57G also weakens ELC binding to myosin but to a lesser extent than E143K since its K_D_ for lever-arm binding is closer to the native ELC (11). Higher duty ratio overcompensates a potentially lower unitary force in A57G giving increased average power. Skinned papillary muscle fibers from E143K transgenic mice showed increased maximal force compared to control (10) that contradicts the lower in vitro isometric force for E143K compared to control (**Table 1**). We hypothesize that the higher duty ratio E143K mutant in the intact muscle recruits some of the normally super-relaxed myosins into the active cycle. These new recruits increase the total number of active myosins giving an overall more robust contraction. In vitro, there is probably no effective super-relaxed myosin pool available from which to recruit hence WT and mutant myosin have identical active myosin totals. For an equal number of myosins, higher duty ratio undercompensates lower unitary force in E143K and in contrast to A57G. It also suggests that secondary remodeling in HCM reflects the inadvertent excess cross-bridge recruitment that might be reversed by a therapy to lower V_max_.

The HCM mutants A57G and E143K up and down regulate power, respectively, complicating a presumption that HCM associates with “hypercontractilty”. Hypercontractility is a general term used to indicate enhancement to one or more motor characteristics including: V_max_, active tension, contractile kinetics, motility velocity, power, and probably others. The present work suggests correlation between single myosin mechanical characteristics and phenotype is nuanced and more likely to be solved when a larger number of the diseased linked mutants are studied using the Qdot assay.

## CONCLUSION

The WT, A57G, and E143K α-myosin variants down-shift their step-size under load like their native in vivo counterpart in embryonic zebrafish. Step-size down-shifting is accomplished by affecting step-size or step-frequency of the unitary steps. The WT and mutant myosins differ substantially in their mechanical characteristics with A57G and E143K consuming more ATP than WT while under load and with A57G up regulating and E143K down regulating power compared to WT. The E143K’s inability to produce more power despite more ATP consumption is probably from a less rigid lever-arm that lowers unitary force.

In vitro results from the loaded Qdot assay presented here and earlier in vivo results (6) indicate varied combination of step-sizes for specific loaded conditions does not necessarily require hierarchical mechanical properties acquired by motor integration into whole muscle. The evident and quantifiable complementarity between in vivo and in vitro systems gives confidence the latter reliably replicates core aspects of the former thus simplifying our future efforts to broadly mechanically characterize inheritable cardiac diseases implicating the cardiac motor. A wide survey of single myosin mechanical characteristics for native and disease linked mutations using the high throughput Qdot assay promises to facilitate therapeutics development with objective intelligent designs suggested from machine learning.

## ACKNOWLEDGEMENT

This work was supported by NIH grants R01AR049277 (to TPB) and R01HL123255 (to DS-C) and by the Mayo Foundation. We thank Katalin Ajtai for her scientific insights and critical evaluation of the manuscript.

